# A Rice Dual-localized Pentatricopeptide Repeat Protein is involved in Organellar RNA Editing with MORFs

**DOI:** 10.1101/223115

**Authors:** Haijun Xiao, Yanghong Xu, Chenzi Ni, Qiannan Zhang, Feiya Zhong, Jishuai Huang, Yingguo Zhu, Jun Hu

## Abstract

Flowering plants engage in diverse RNA editing events in mitochondrion and chloroplast on post-transcriptional process. Although several PPRs and MORFs were identified as RNA editing factors, the underlying mechanism of PPRs and the cooperation among them are still obscure. Here, we identified a rice dual-localized PPR mutant *Ospgl1*. Loss-of-function of *OsPGL1* resulted in defect of chloroplast RNA editing at *ndhD-878* and mitochondrial RNA editing at ccmFc-543, which can be restored via complementary validation. Despite the synonymous editing on ccmFc-543, loss of editing at *ndhD-878* caused failure of conversion from serine to leucine, leading to the dysfunction of chloroplast and defective in photosynthetic complex, further studies demonstrated OsPGL1 directly bound to both two transcripts. The interaction between three MORFs (MORF2/8/9) and OsPGL1 were confirmed *in vitro* and *in vivo*, implied OsPGL1 functioned on RNA editing via an editosome. It also suggested MORFs assisted and contributed to the flexible PPR-RNA recognition model during RNA editing through the cooperation with PPRs. These results provide new insight into the relationship between RNA editing and plant development on chloroplast.

**Highlight:** We firstly characterized a dual-localized PPR protein which is required for RNA editing in mitochondrion and chloroplast simultaneously. OsPGL1 binds to two distinguish target transcripts directly and cooperated with MORFs.

## Introduction

RNA editing was broadly defined as a post-transcriptional process that changes the sequence of an RNA molecule from that of its DNA master (Covello and Gray, 1989). RNA editing is widely spread in eukaryotic cell and highlighted in plant mitochondria and chloroplasts. In animals, RNA editing was mediated by two mechanisms, one is C-to-U editing by apolipoprotein B mRNA editing enzyme, catalytic (APOBEC) and another is adenosine (A)-to-inosine (I) editing by adenosine deaminase acting on RNA (ADAR) (Kim *et al.*, 1994; Mehta and Driscoll, 2002; Melcher *et al.*, 1996; Teng *et al.*, 1993). In plants, RNA editing includes C-to-U, U-to-C and A-to-I conversion (Takenaka *et al.*, 2013b). The majority of editing in plants occurs in mitochondrial and plastid transcripts, however, A-to-I editing also occurs in cytosolic tRNAs (Chateigner-Boutin and Small, 2010). RNA editing from C-to-U is the most frequent editing events in plants, in Arabidopsis and rice, 525 and 491 C-to-U editing sites in mitochondria, 34 and 21 C-to-U editing sites in chloroplast has been identified in previous study (Chateigner-Boutin and Small, 2010). In human, the APOBEC3 proteins can deaminate cytidines to uridines in single-stranded DNA (ssDNA) (McDougall *et al.*, 2011). Recently, the fusion of APOBEC3 with catalytically dead Cas9 (dCas9) or other Cas9 variants in CRISPR system accomplished the genomic editing of single bases in mammalian, yeasts and plants (Hess *et al.*, 2016; Kim *et al.*, 2017; Lu and Zhu, 2017; Ma *et al.*, 2016; Zong *et al.*, 2017). While in plants, the deaminase activity still need to be confirmed and elucidated. Researchers proposed RNA editing activity is governed by RNA-binding pentatricopeptide repeat (PPR) proteins, and DYW-subclass PPR proteins were considered as mainly RNA editing factors depend on its similarity with cytidine deaminases of the DYW motif (Salone *et al.*, 2007)

To date, several non-PPR editing factors, such as RNA editing factor interacting proteins (RIPs)/multiple organellar RNA editing factors (MORFs), organelle RNA recognition motif (ORRM) proteins, organelle zinc-finger (OZ) proteins, and protoporphyrinogen oxidase 1 (PPO1) have been identified as components of the plant RNA apparatus (Sun *et al.*, 2016). There were nine MORF proteins in Arabidopsis and five members in rice. MORF8/RIP1 was identified as an interacting factor of RARE1, a PPR protein required for RNA editing in Arabidopsis chloroplast (Bentolila *et al.*, 2012). MORF2 and MORF9 were both targeted exclusively in plastids and can affect most of RNA editing sites in chloroplast in Arabidopsis (Takenaka *et al.*, 2012). MORF2/8/9 can interact with ORRM6, which is involved in RNA editing of psbF-C77 and accD-C794 in Arabidopsis (Hackett *et al.*, 2017). Furthermore, MORFs also interact with each other and form a heterodimeric or homodimeric complex, suggesting a more complicated regulation mechanism in plants (Takenaka *et al.*, 2012).

The PPR protein family was characterized by the degenerate motifs of 35 amino acids arranged as tandem repeats of 2–25 such elements, settled in a pair of antiparallel double alpha-helices, helices A and B (Small and Peeters, 2000; Yin *et al.*, 2013). PPR proteins have been found to be in eukaryotic genomes and greatly expanded in the plants, study showed more than 400 members harbored into land plants (Cheng *et al.*, 2016). PPR protein can be divided into two sub-families: P and PLS subfamilies, the P sub-families proteins contain tandem arrays of canonical 35-amino-acid (P) PPR repeats, whereas PLS sub-families characterized by triplets of P, L (i.e.35 to 36 amino acids in length), and S (i.e.31 amino acids) motifs (Lurin *et al.*, 2004). Many post-transcriptional processes in these organelles were relevant to PPR proteins, including RNA editing, splicing, cleavage, RNA stability, and translation (Schmitz-Linneweber and Small, 2008). Most PPR proteins are targeted either mitochondria or chloroplasts, but few of them are dual-localization.

Here, we addressed a novel dual-localized PPR protein OsPGL1 in rice, which is required for RNA editing at two different *cis* elements. Loss of function of *OsPGL1* caused a pale green leaves phenotype, resulting from defective photosynthetic complex in chloroplast development. Loss of editing at *ndhD-878* caused failure of conversion from serine to leucine, which an extremely conserved amino acid in plants. Further investigation showed OsPGL1 functions with MORF2/8/9 and directly binds to *ndhD* and *ccmFc* via its 9 PPR motifs. These results confirmed PPR proteins and MORFs are required for RNA editing, and function together via a complex editosome.

## Materials and Methods

### Plasmid construction and transformation

Two target sites at the 20 bp upstream of protospacer-adjacent motif sequence (PAM) according to the recognition principle of CRISPR/Cas9 were designed and analyzed the specificity by CAS-OFFinder (http://www.rgenome.net/cas-offinder) (Table S1). The target sequence joint was linked into gRNA-U3 and gRNA-U6 vector, respectively followed by two rounds of nest-PCR. PCR products were subsequently linked to CRISPR/Cas9 vector. The construction was identified through PCR and sequencing. Calli derived from ZhongHua 11 (*Oryza sativa. L. Japonica*) were used for *Agrobacterium*-mediated transformation. WT and CRISPR/Cas9 knockout lines were grown in a paddy field and greenhouse in Wuhan, China under proper management.

### Scanning electron microscopy and transmission electron microscopy assay

For scanning electron microscopy assay, samples were prepared as described previously (Zhou *et al.*, 2011). Rice young leaves were cut into small section with a razor and immediately placed in 70% ethanol, 5% acetic acid, and 4% formaldehyde for 18 h. Samples were critical point dried, sputter coated with gold in an E-100 ion sputter, and observed with a scanning electron microscopy (Hitachi S-3000N, Japan).

For transmission electron microscopy, samples were fixed in 2.5% (w/v) paraformaldehyde and 0.25% glutaraldehyde in 0.2 N sodium phosphate buffer for 2–4 h at 4 °C, pH 7.0, and were then post-fixed in 1% OsO_4_ in PBS, pH 7.4. Following ethanol dehydration, samples were embedded in acrylic resin. Ultrathin sections (50 to 70 nm) were double stained with 2% (w/v) uranyl acetate and 2.6% (w/v) lead citrate aqueous solution and examined with a transmission electron microscope at 200 kV (Tecnai G2 20 Twin, FEI, Netherlands).

### RNA Extraction and qRT-PCR

Total RNA was extracted with 1 mL Trizol reagent according to the manufacturer’s instructions (Invitrogen). After isopropanol precipitation, the RNA was resuspended in 30 μl RNase-free water and treated with RNase-free DNase I (New England Biolabs). First-strand cDNA was reverse transcribed using random primers (Primers were listed in Table S2). Ubiquitin was detected as control for gene expression.

### Analysis of RNA editing

For RNA editing analysis in the wild type and the *Ospgl1*, total RNAs were isolated from the young leaves using the Trizol reagent as described before (Hu *et al.*, 2012). RNA was treated with RNase-free DNase I (New England Biolabs), and confirmed by PCR. Then, the RNAs were reverse transcribed with random primers and the high-fidelity reverse transcriptase SuperScript III (Invitrogen). Primers were designed to cover all 491 mitochondrial editing sites and 21 chloroplast editing sites (Table S2). The RT-PCR products were sequenced directly.

### Subcellular localization of OsPGL1

For transient expression in rice protoplast, 188 amino acid of the OsPGL1 N-terminal were cloned into HBT-sGFP driven by the cauliflower mosaic virus 35S promoter to construct the 35S:OsPGL1^N1-188^:sGFP fusion protein. Protoplast preparation and transformation procedures were as previously described (Yu *et al.*, 2014). MitoTracker Red (Invitrogen) was used as a mitochondrial specific dye.

### RNA electrophoresis mobility shift assays (REMSA)

The corresponding cDNA fragments of *OsPGL1* was amplified with specific primers (Table S2), and cloned into the pGEX-6p-1 vector to create the fusion protein GST-OsPGL1. Two RNA probes (probe 1 and probe 2) and negative control probe (probe C) containing the target editing site were synthesized and labeled with biotin at the 3’ end by GenScript (Nanjing, China). For REMSA, the recombinant protein was incubated with RNA probe in a 20 μl reaction mixture including 10 μl of 2 × binding buffer (100 mM Na phosphate, pH 7.5, 10 units RNasin, 0.1 mg/mL BSA, 10 mM DTT, 2.5 mg/ mL heparin, and 300 mM NaCl). The mixture was incubated at 25 °C for 30 min, followed with separation by 5% native PAGE in 0.5×TBE buffer and transferred onto the nylon membrane (Roche). For the competitive REMSA, the gradually increased concentration of the unlabeled probe was added into the reaction mixture followed the procedure described above.

### Complementation of *Ospgl1* mutants

For complementation of the *Ospgl1* mutant, a full-length (1815bp) cDNA fragment was constructed into pCAMBIA-2300 vector driven by CMV-35S promoter and transformed into *Ospgl1-1* mutant background by agrobacterium-mediated method. Independent transgenic lines were obtained and planted in Wuhan, China.

### Yeast two-hybrid assays

The full-length cDNA of *OsPGL1* and *MORFs* were cloned into pGBKT7 and pGADT7 vector. The constructs were co-transformed into yeast (AH109 strain) in pairs according to previous study (Hu *et al.*, 2012).

### GST pull-down assays

The purified recombinant proteins (GST tag, Trx-His tag, GST-OsPGL1, Trx-MORF2-His, Trx-MORF8-His and Trx-MORF9-His) were dialyzed against phosphate-buffered saline (PBS; 137 mM NaCl, 2.7 mM KCl, 10 mM Na_2_HPO_4_, 2 mM KH_2_PO_4_) for 24 h and quantified using thebicinchoninic acid (BCA) method. The recombinant protein GST-OsPGL1 was incubated with glutathione sepharose for 1 h on ice and washed with five volumes of PBS for five times. Trx-His, Trx-MORFs-His proteins were subsequently added to detect the interaction. The binding proteins were washed with five volumes of PBS for five times, eluted with glutathione reductase and separated by 10% SDS-PAGE. Products were transferred onto a polyvinylidene fluoride (PVDF) membrane (BioRad), and investigated with antibodies against GST and His respectively.

### BiFC assays

For bimolecular fluorescence complementation (BiFC) analysis, the full-length cDNA of OsPGL1 without the stop codon was fused to the C-terminal fragment of yellow fluorescent protein (YFP) in pUC-SPYCE (C-terminal), MORF2, MORF8 and MORF9 were fused to the N-terminal fragment of YFP in pUC-SPYNE (N-terminal). The two vectors were co-transformed into rice protoplasts in pairs and observed by bright-field and fluorescent microscopy using a Leica DM4000B microscopy (Hu *et al.*, 2012).

### Co-immunoprecipitation analysis

Total proteins from transgenic plants fused with FLAG (UBI: OsPGL1-FLAG) and GFP (35S: MORFs-GFP) were extracted with extraction buffer (100 mM Tris-HCl, 200mM NaCl, 5 mM EGTA, 5 mM EDTA, 10 mM DTT, 0.6% TritonX-100, 1 mM PMSF, pH 8.0) and incubated with 2μg anti-GFP antibody overnight at 4°C, 100 μl of protein A-Sepharose beads was added and incubated for a further 3 to 4 h. Immunoprecipitates were washed for five times with co-immunoprecipitation buffer, (150mM NaCl, 20mM Tris-HCl, 1mM EDTA, 0.2% NP-40, 1 mM PMSF, pH 7.4) and loaded in 6×SDS loading buffer by denaturing for 10 min. The proteins were separated on a 10% SD-SPAGE gel and detected by immunoblotting with anti-FLAG and anti-GFP antibodies.

### Immunoblot analysis

Total proteins were extracted from young leaves and quantified with the BCA protein assay kit (Thermo scientific). 10 μg total proteins were separated by SDS-PAGE, transferred onto a polyvinylidene difluoride (PVDF) membrane (Bio-rad) and incubated with various primary antibodies against NdhD (Beijing Protein Innovation, China), PsaA, PsbA, PetA, AtpA, Lhca2 and Lhcb2 (Agrisera). Actin was used as a reference antibody. Detection was carried out by the ECL western blotting detection reagents (Bio-rad).

### Blue-native PAGE

The equivalent of 500 μg of total mitochondrial proteins from WT and *Ospgl1-1* mutant were treated and loaded on BN-PAGE according to the previous protocol (Liu *et al.*, 2012). The gel was firstly stained with coomassie brilliant blue. For immunoblotting, the gel was transferred to the PVDF membranes, antibodies against Cytc1 were used to detect the accumulation of complex III.

### Accession numbers

Sequence for rice OsPGL1, chloroplast NdhD, and mitochondrial gene CcmFc can be found in the GenBank database under accession numbers XP_015618645.1 (OsPGL1), NP_039444 (NdhD), YP_002000589 (CcmFc), respectively.

## Results

### Phenotypic characterization of the *Ospgl1* mutant

We used Clustered Regularly Interspaced Short Palindromic Repeat (CRISPR) system to generate several PPRs mutant under the background of the ZhongHua 11 (ZH11, *Oryza satia. L. Japonica*) and obtained two independent transgenic knocked-out lines of LOC_Os12g06650. Further genome DNA sequencing revealed a 41 bp deletion from 288 to 328 and a 1 bp deletion at 530 in this PPR gene (Fig. 1A-C). Both two lines exhibited pale green leaves at all vegetative stages (Fig. 1D-F), thus we named this gene as *pale green leaves 1* (*Ospgl1*) in this study. And we defined these two lines as *Ospgl1-1* and *Ospgl1-2*, respectively. The content of chlorophyll was drastically reduced than those in WT plants (Fig. 1G). This phenotype performed more pronounced in paddy fields (Fig. S1A). Despite of the distinct development on vegetative stage, the plant height, tiller number and seed setting were rarely impaired (Fig. 1H-J). To confirm the phenotype of *Ospgl1* knockout mutants, we also generated the transgenic RNAi lines. Results showed that suppression of *OsPGL1* expression recapitulated the knockout mutants phenotype (Fig. S2).

**Figure 1.**
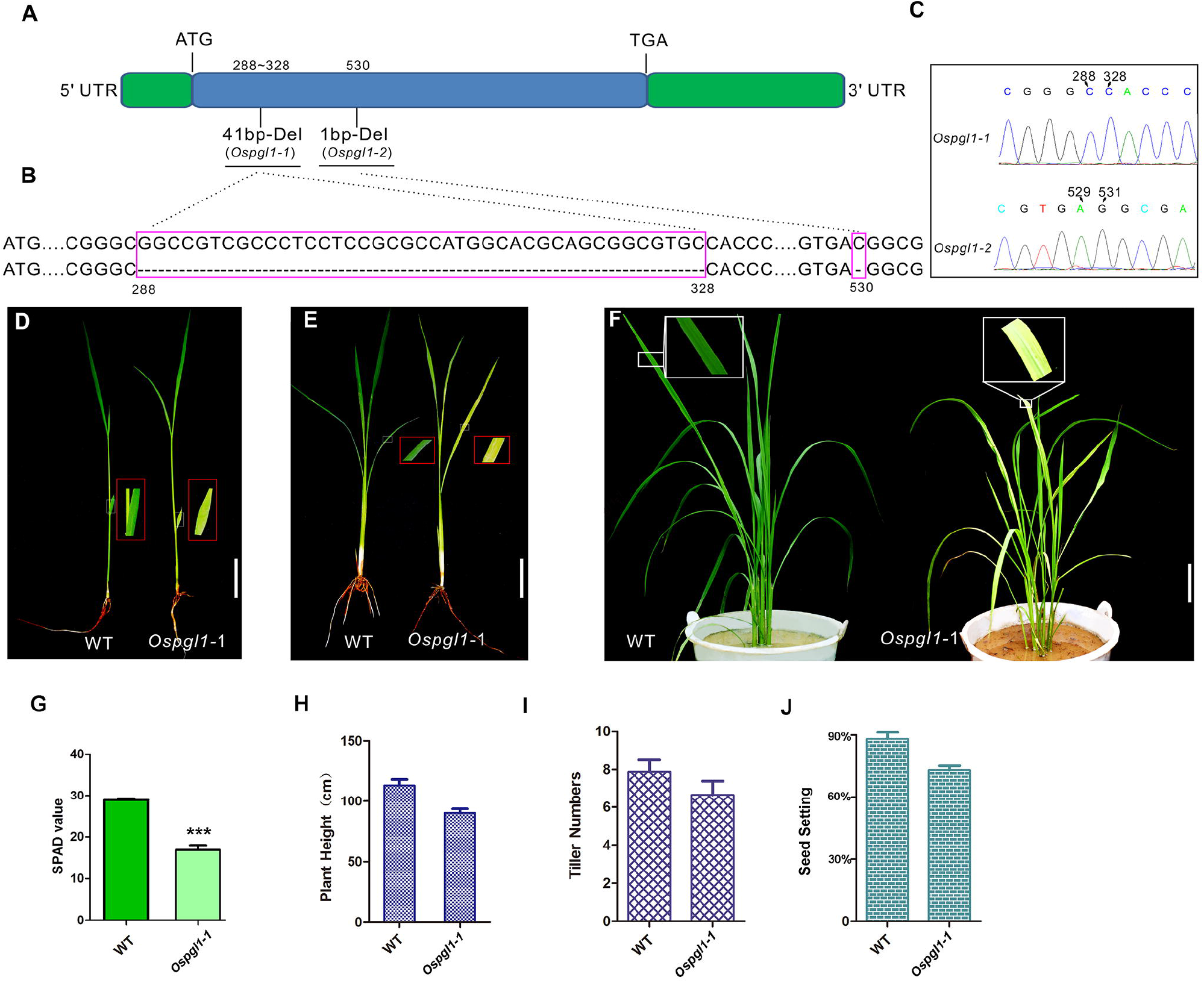
Mutant and phenotypic characterization of *Ospgl1*. **(A)** Schematic drawing of the intronless gene *OsPGL1* attaching the position of the deletion in *Ospgl1*. **(B)** Alignment of WT and mutant DNA sequences around the target sites highlighting the deletion of 41bp in *Ospgl1-1* and 1bp deletion in *Ospgl1-2* marked with a pink box. **(C)** Sequencing results of *Ospgl1-1* and *Ospgl1-2* heterozygotes. **(D)** Comparison of leaves from *Ospgl1-1* and WT plant at the two-leaf stage, the red box shows magnified view, bar = 2cm. **(E)** Comparison of leaves from *Ospgl1-1* and WT plant at the three-leaf stage, the red box shows magnified view, bar = 2cm. **(F)** Comparison of leaves from *Ospgl1-1* and WT plant at the tillering stage, the red box shows magnified view, bar = 5cm. **(G)** Comparison of the chlorophyll content in *Ospgl1-1* and WT. Bars represent mean ± SD from three independent biological replicates. Asterisks indicate statistically significant differences compared with the WT (Student’s t-test: *** P< 0.001). **(H)** Comparison of the plant height of *Ospgl1-1* and WT plant. **(I)** Comparison of the tiller number of *Ospgl1-1* and WT plant. **(J)** Comparison of the seed setting of *Ospgl1-1* and WT plant.

To dissect the morphologic details of the leaf, we performed scanning electron microscopy (SEM) and transmission electron microscopy (TEM) examinations. SEM results showed the leaf is complete and intact in WT (Fig. 2A), while some cracked holes were distributed in the leaf of *Ospgl1-1* (Fig. 2F). Furthermore, TEM assays showed the starch granular stacks in WT plant were well-balanced (Fig. 2B), while the starch granular stacks in *Ospgl1-1* were reduced and incompact (Fig. 2G). When we looked insight into the structure of thylakoid, results showed the density of well-organized thylakoid were not observed in *Ospgl1-1*, which exhibited more hollow structures (Fig. 2C, 2H). In spite of the fact that chloroplasts were impaired, the ultrastructure of the mitochondria was undistinguishable between WT and *Ospgl1-1* (Fig. 2D, 2E and 2I, 2J). These observations revealed that the *Ospgl1* mutation primarily affects the development and morphology of chloroplasts rather than mitochondria.

**Figure 2.**
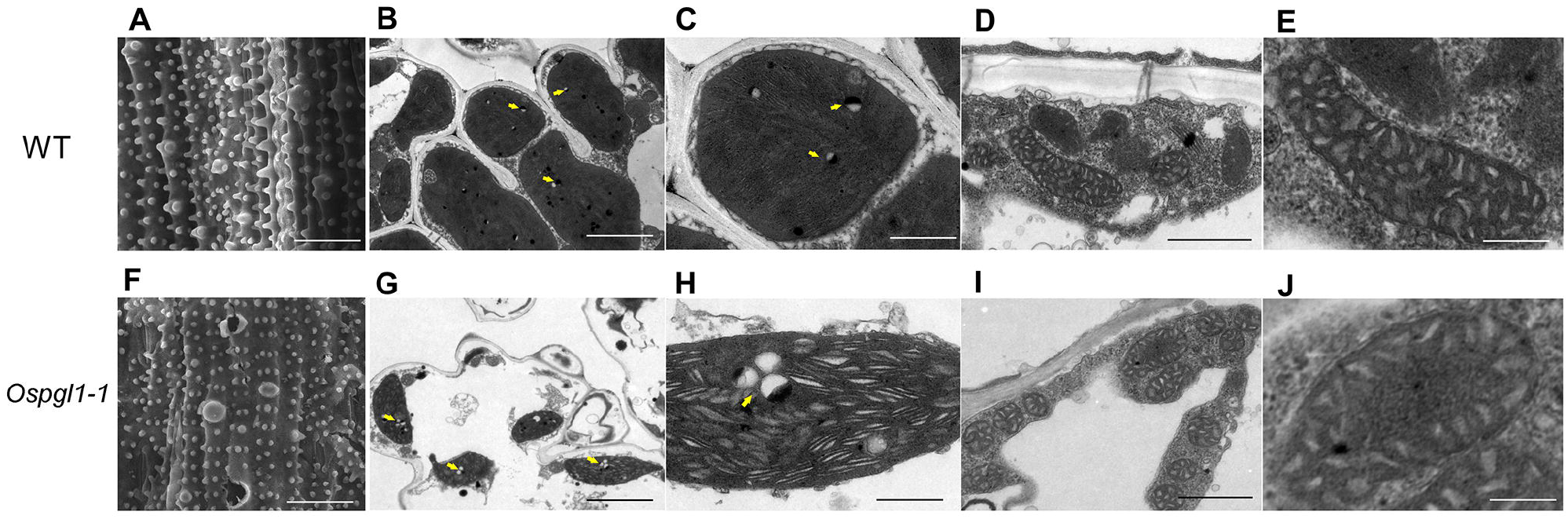
Scanning electron microscopy and transmission electron microscopy examination of leaves at tillering stage of WT and *Ospgl1-1*. **(A)** Scanning electron microscopy image of WT leaf, bar = 20 μm **(B)** Transmission electron microscopy image of WT chloroplasts, bar = 2μm **(C)** A higher magnification image of WT chloroplast from **(B)**, bar = 0.3μm **(D)** Transmission electron microscopy image of WT mitochondria, bar = 0.5μm **(E)** A higher magnification image of WT mitochondrion from **(D)**, bar = 0.15μm **(F)** Scanning electron microscopy image of mutant leaf, the image shows a defective Leaf surface morphology with cracked, broken holes at the pale parts of *Ospgl1-1*, bar = 20 μm **(G)** Transmission electron microscopy image of chloroplasts in pale parts of *Ospgl1-1* mutant leaf, the image shows incompact granal stacks and lacked well-structured thylakoid, bar = 1μm **(H)** A higher magnification image of chloroplast in pale parts of *Ospgl1-1* mutant leaf from **(G)**, bar = 0.3μm **(I)** Transmission electron microscopy image of mitochondria in *Ospgl1-1* leaf, bar = 0.3μm **(J)** A higher magnification image of *Ospgl1-1* mitochondrion from **(I)**, bar = 0.15μm

### *OsPGL1* encodes a DYW-motif containing protein

*OsPGL1* encodes a putative PPR protein consisting of 605 amino acids without intron. Motif prediction analysis by Pfam (http://pfam.xfam.org/) revealed that OsPGL1 consists of 9 PPR motifs. 3 S motifs, 3 P motifs and 3 L motifs present staggered arrangement ( Fig. S3A, 3B). The C-terminal region from residues 392 to 605 shows the consensus sequences of the extension domains (E, E+, and DYW domains) (Fig. S4). This data indicated that OsPGL1 belongs to the typical DYW type of PLS subfamily. Alignment of *OsPGL1* with its orthologs in various plants showed 75%, 75%, 46%, 51%, 51%, 47% similarity with *Zea mays* (GRMZM2G001466), *Sorghum bicolor* (Sb08g003980), *Arabidopsis thaliana* (AT4G15720,REME2), *Theobroma cacao* (XP_017974392), *Glycine max* (XP_003529581), and *Brassica napus* (BnaC07g33170D) Fig. S4). Previous study reported that REME2 is involved in RNA editing of rps3-1534 and rps4-175 in mitochondria (Bentolila *et al.*, 2014). The high similarity among OsPGL1, GRMZM2G001466 and Sb08g003980 implied the function of this PPR gene might be conserved in monocots. Based on the PPR-RNA recognition mode, we analyzed the conservation of the 1 ‘ and 6 amino acid of each motif, which is essential for RNA recognition. Results showed these candidate orthologs could be divided into two subgroup, monocots and dicots, which suggested the functional conservation respectively (Fig. S5). It also implied the distinct function between OsPGL1 and AtREME2.

### Expression pattern of *OsPGL1*

To get insight into the expression pattern of *OsPGL1* in WT, RT-PCR and quantitative RT-PCR (qRT-PCR) were employed to detect various tissues. RT-PCR Results showed that *OsPGL1* was constitutively expressed in both vegetative and reproductive tissues, including root, stem, leaf, and panicle (Fig. 3A). Moreover, qRT-PCR data showed the expression level in root, stem and panicle, but preferentially accumulated in fresh leaf (Fig. 3B). These data are consistent with the results of the leaf phenotype, and slightly effects on plant height and seed setting.

**Figure 3.**
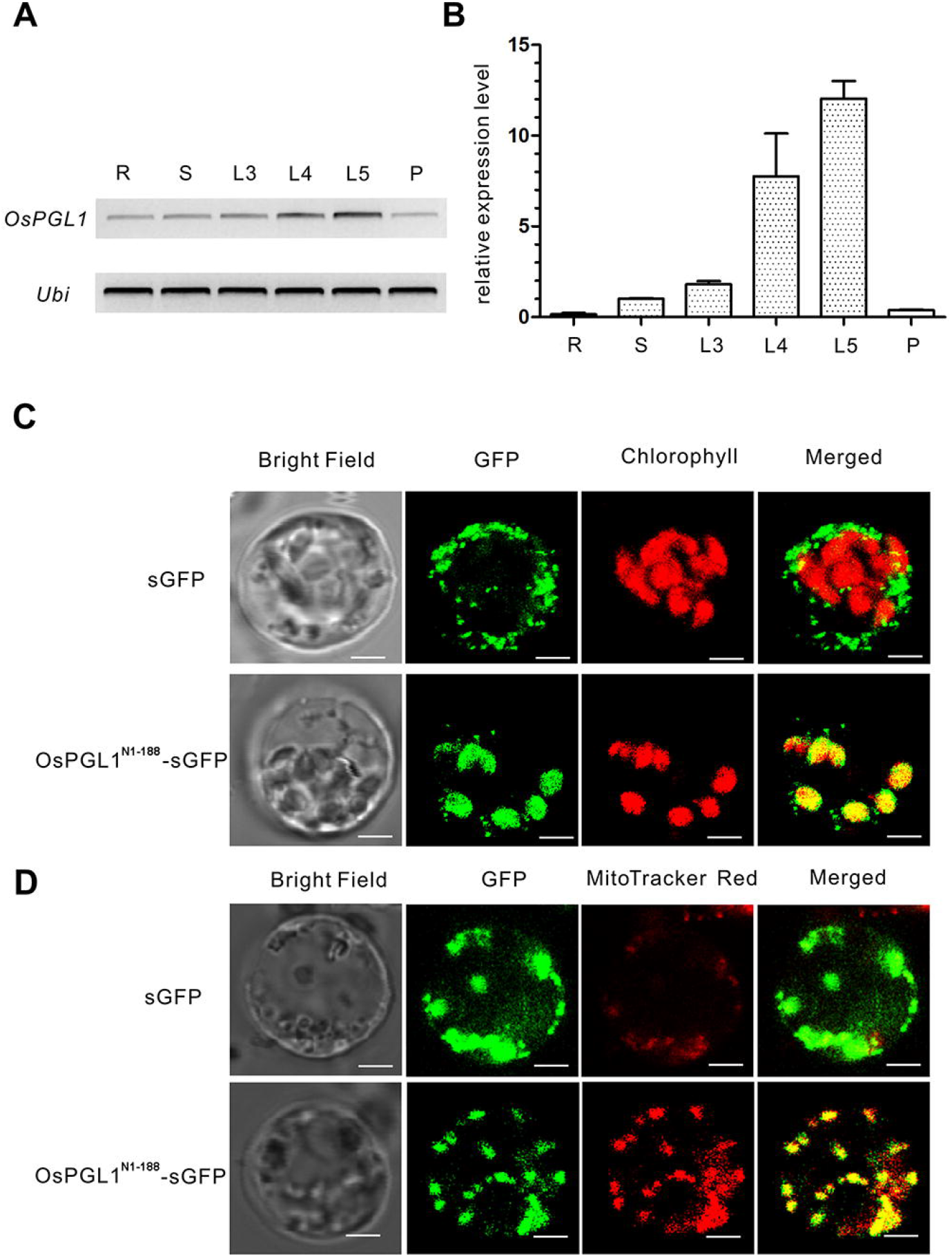
Expression and subcellular localization analysis of *OsPGL1*. **(A)** RT-PCR examination of *OsPGL1* in different tissues of WT. R, Root; S, Steam; L3, three-leaf stage; L4, four-leaf stage; L5, three-leaf stage, five-leaf stage; P, panicle. **(B)** Real time RT-PCR examination of *OsPGL1* in different tissues of WT. Error bars represent the SD. **(C)** Transient expression of 35S:sGFP (top) and 35S:OsPGL1^N1-188^-sGFP (bottom) in rice protoplast. Bar = 5 μm. **(D)** Transient expression of 35S:sGFP (top) and 35S:OsPGL1^N1-188^-sGFP (bottom) in rice protoplast. MitoTracker Red were used for mitochondria indicator. Bar = 5 μm.

### OsPGL1 is a novel dual-localized PPR protein

Plenty of reports confirmed that most PPR proteins are targeted to plastids, chloroplast or mitochondrion. With bioinformatic analysis of TargetP (http://www.cbs.dtu.dk/services/TargetP/), *OsPGL1* was predicted to localize into chloroplast, and also into mitochondrion with a low degree of confidence. To make sure of the subcellular localization of OsPGL1, the N-terminal region (amino acids from 1 to 188) were fused with green flourecent protein (sGFP), driven by CaMV35S promoter and transiently expressed in rice protoplast. Results showed the GFP signals could overlap the red auto-fluorescent signals of chlorophyll (Fig. 3C). Interestingly, there were still some spots could not overlap the signals from chloroplast, implied OsPGL1 might target into mitochondria. Subsequently, mitotracker red were used to indicate mitochondrion, which signals were also overlapped with the signals of OsPGL1-GFP (Fig. 3D). Therefore, OsPGL1 is a novel dual-localized PPR protein in rice, both targets into mitochondria and chloroplasts.

### OsPGL1 is involved in C-to-U RNA editing of *ndhD* and *ccmFc* transcripts

Studies showed PPR proteins were involved in the RNA editing of one or several editing sites, especially in DYW subfamily (Sun *et al.*, 2016). Consideration of the dual-localization of OsPGL1, we checked all 491 editing sites in mitochondria and 21 editing sites in chloroplasts respectively by RT-PCR. Sequencing results revealed that the C-to-U editing efficiency of *ndhD-878* and *ccmFc-543* dramatically decreased to zero in both two mutants expect for 8% at *ccmFc-543* in *Ospgl1-2*, while *ndhD-878* and *ccmFc-543* were completed edited in WT (Fig. 4A). Consequently, abolishment of editing in *ndhD-878* recovered the codon from UUA to UCA, which resulted in the amino acid substitution from leucine to serine. The loss of editing in another site ccmFc-543, a mitochondrial RNA editing site converted the codon from GUU to GUC, due to the degeneracy of codon, the change of the nucleotide at this position does not cause any amino acid alteration, consistent with the results of no effects on mitochondrion in *Ospgl1* (Fig. 2I, 2J).

**Figure 4.**
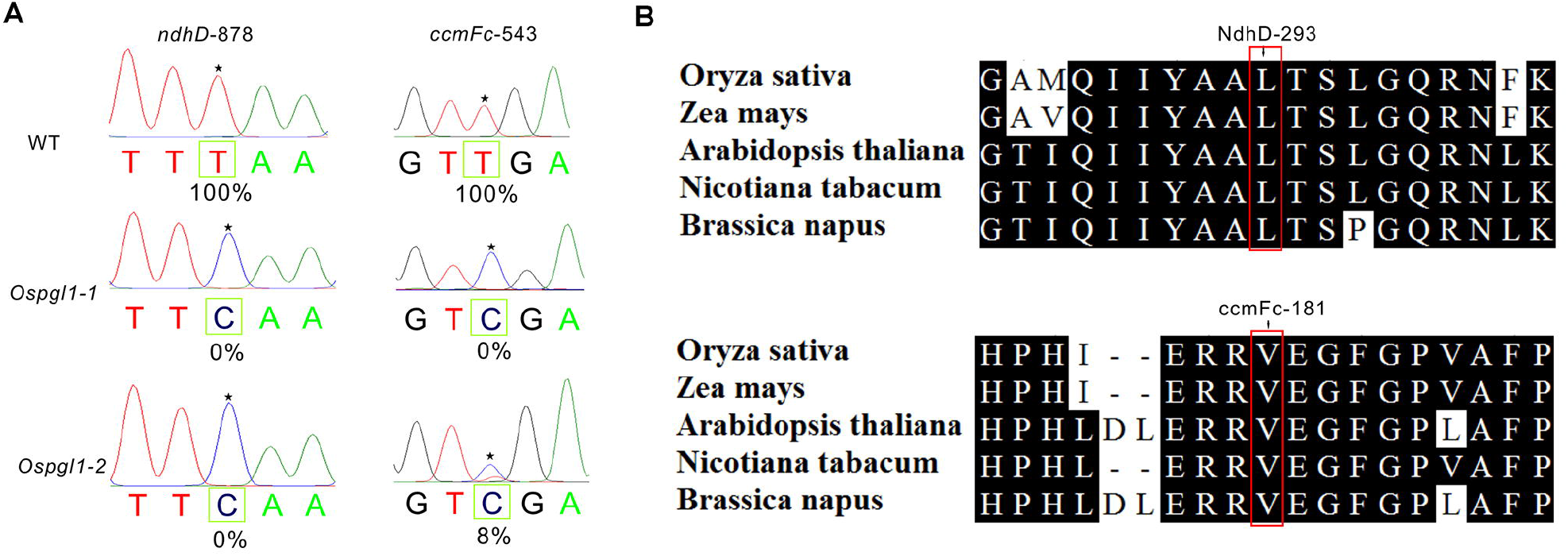
Organelle RNA editing analysis in different species. **(A)** RNA editing analysis of the *ndhD-878* and ccmFc-543 sites from WT and *Ospgl1-1* leaves. The black stars marks the editing site, the editing efficiency was presented under the target sites. **(B)** Alignment of the orthologous NdhD and CcmFc amino acids sequences in five different species around the corresponding affected residues. Numbers and the red rectangular box indicate the converted amino acid.

To investigate the evolutionary conservation of the alteration from serine to leucine at the 293 position of NdhD protein sequence and valine at the 181 position of CcmFc protein sequence, the chloroplast NdhD orthologs and mitochondrial CcmFc orthologs from five representative species (*Oryza sativa, zea mays, Arabidopsis thaliana, Nicotiana tabacum and Brassica napus*) in plants were analyzed. Results showed these two residues are extremely conserved in the five tested species including monocots and dicots, implied that the leucine of NdhD and valine of CcmFc could important for plant development (Fig. 4B).

### OsPGL1 can bind to both *ndhD* and *ccmFc* transcripts

To test the RNA binding activity of OsPGL1, we expressed recombinant OsPGL1 for further RNA electrophoresis mobility shift assays (REMSA). 9 PPR motifs from residues 46 to 605 was fused with a glutathione S-transferase (GST) tag for expression in *Escherichia coli*. The recombinant protein (GST-OsPGL1^46-605^) was analyzed by western blot with anti-GST antibody to confirm the high purity (Fig. S7b). Subsequently, the recombinant protein was dialyzed to remove the contamination of RNase for REMSA and further quantified. The *ndhD* and *ccmFc* probes include 35 nucleotides surrounding the target editing sites were prepared and designated as probe 1 and probe 2. Probe C (nad3-155) was used as a negative control probe which has been reported as a specific target of another rice PPR protein (manuscript is preparing) (Fig. 5A). GST-OsPGL1^46–604^ and GST tag were incubated with the biotin-labeled RNA probes, respectively. Both of the two protein–RNA complexes were detected as a shifted band that migrated more slowly than free RNA probe in the native gel, but no retarded band was observed when incubated with the GST (Fig. 5B, 5C). In addition, as negative control, no retarded band was observed when probe C was incubated with GST-OsPGL1^46–604^. We next performed the competitor assay using non-labeled RNA probe with the same sequence, the binding intensity of the band were decreased accompanied with the increased concentration of competitors (Fig. 5B, 5C). These data validated the OsPGL1 binds to both *ndhD* and *ccmFc* transcripts directed via the PPR motifs. *ndhD* and *ccmFc* derived from chloroplast and mitochondrion also confirmed the dual-localization of OsPGL1.

**Figure 5.**
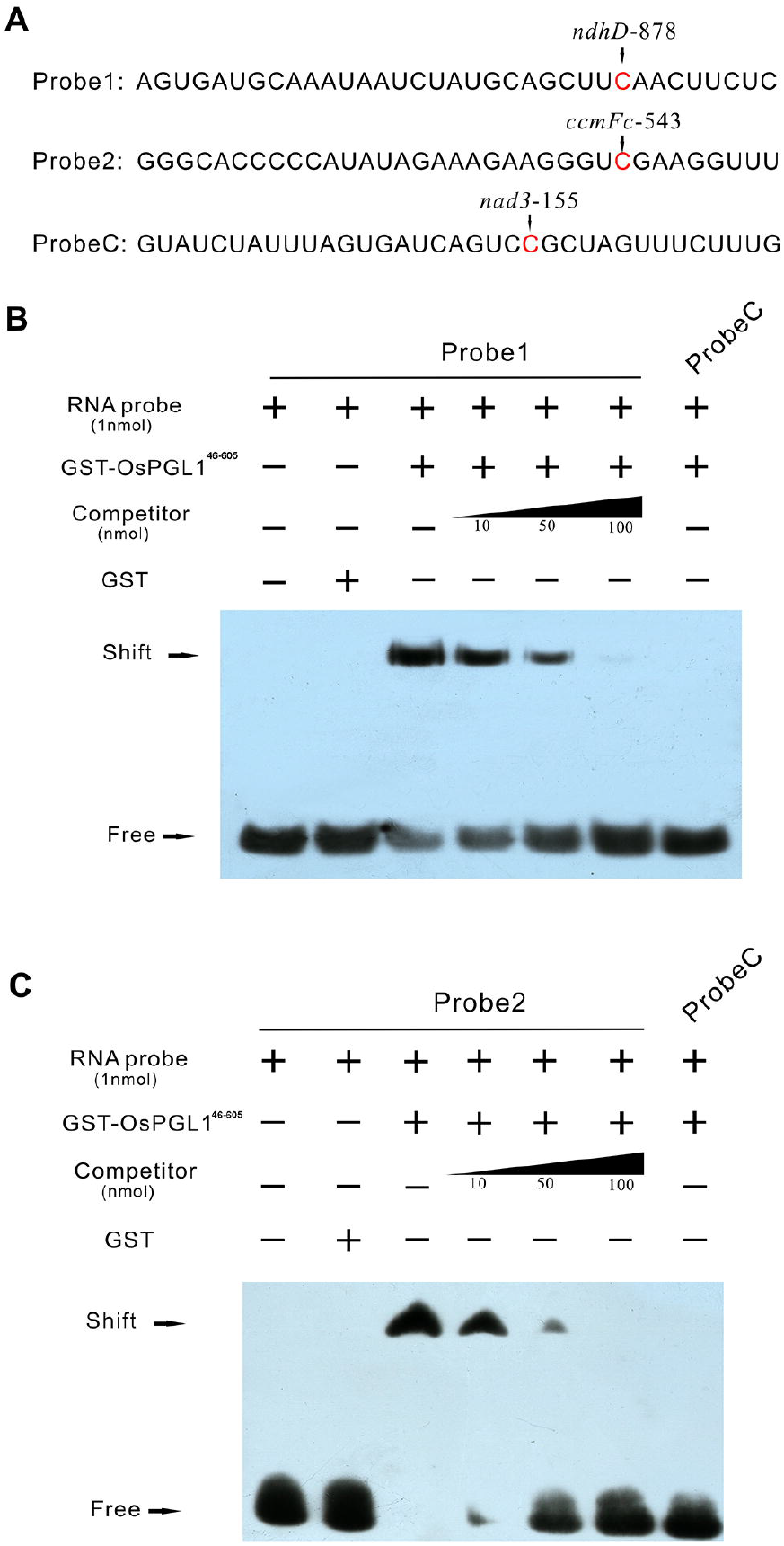
OsPGL1 possesses specific RNA binding activities. **(A)** Schematic sequences of RNA probes. Edited sites were indicated and marked in red letter. Probe C is a negative control in this study. **(B)** RNA electrophoresis mobility shift assays (REMSA) of GST-OsPGL1^46-605^ and GST-tag with RNA probe 1. Unlabeled probe 1 was used as competitor at a range of the concentrations for competitive REMSA. GST-tag and probe C were used as negative control. **(C)** RNA electrophoresis mobility shift assays (REMSA) of GST-OsPGL1^46-605^ and GST-tag with RNA probe 2. Unlabeled probe 2 was used as competitor at a range of the concentrations for competitive REMSA. GST-tag and probe C were used as negative control.

### The transgenic complementation lines rescue the pale green phenotype

To verify whether the pale green phenotype resulted from the dysfunction of *OsPGL1* in deed, we performed the transgenic complementation assay in mutant lines. Full-length coding sequences of *OsPGL1* was constructed into pCAMBIA-2300 vector driven by CaMV-35S promoter and transformed into *Ospgl1-1* mutant by *Agrobacterium-mediated* method. All 12 independent transgenic lines completely rescued the mutant pale green leaves phenotype (Fig. 6A). Furthermore, we checked the RNA editing efficiency of *ndhD-878* and *ccmFc-543* in all 12 independent transgenic complementation lines, results showed 35S:OsPGL1 completely recovered the RNA editing efficiency to 100% same as those in WT (Fig. 6B). Data indicates that the pale green leaves phenotype was in deed caused by loss of RNA editing of *ndhD-878*, which resulted from loss of function of *OsPGL1*.

**Figure 6.**
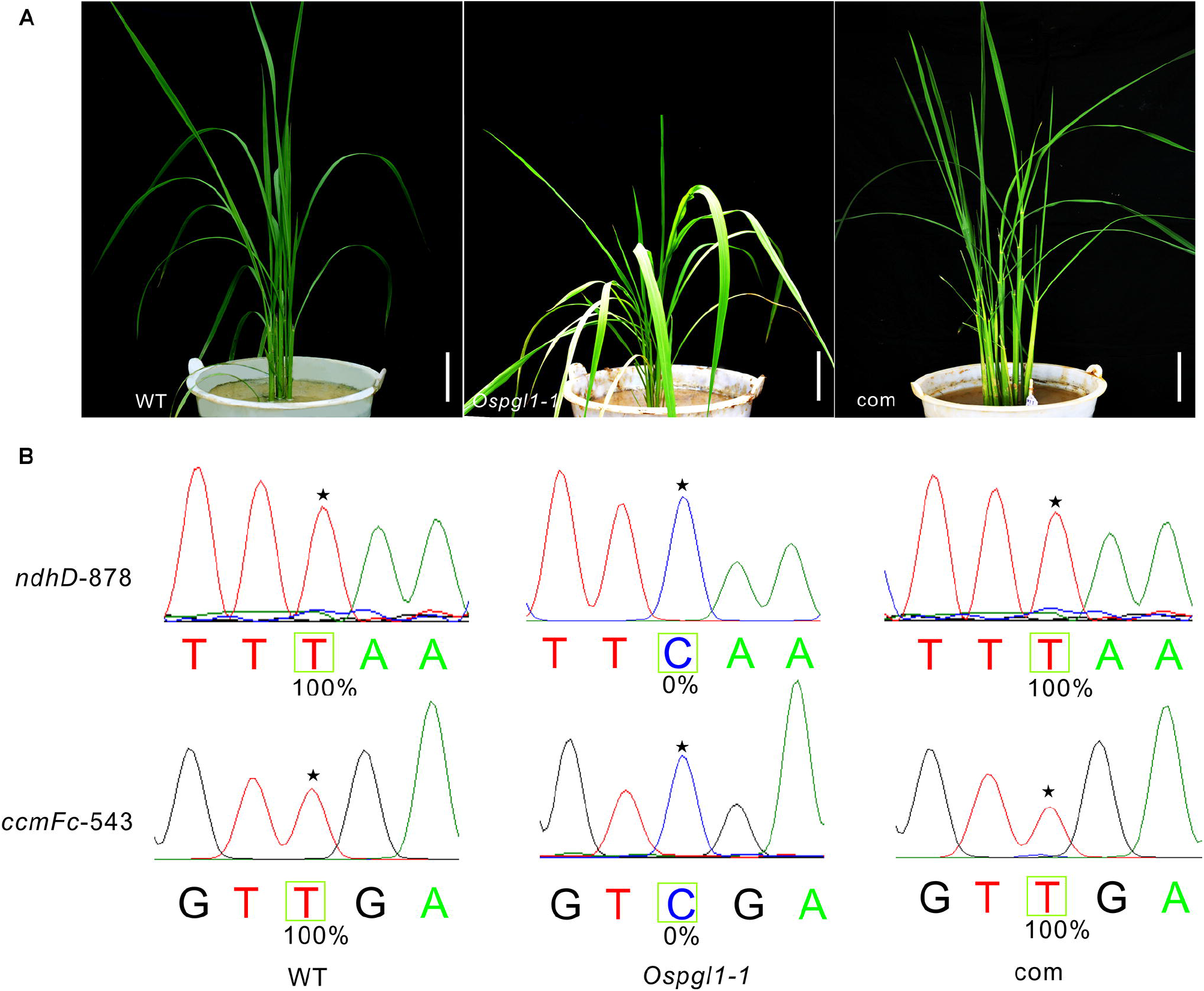
Complementation analysis of *Ospgl1-1*. **(A)** The phenotypes of WT, *Ospgl1-1* and complemented T_0_ plant (com) at the tillering stage. bars = 5cm. **(B)** RNA editing efficiency of *ndhD-878* and *ccmFc-543* comparison among WT, *Ospgl1-1* and complemented T_0_ plant (com), the stars show editing sites.

### OsPGL1 interacts with three MORF proteins *in vitro* and *in vivo*

In addition to PPR proteins, another group of RNA editing factors in plant organelle was identified, the multiple organelle RNA editing factor (MORF) proteins, which has also been termed RNA editing factor interacting protein (RIP) proteins. There were five MORF proteins in rice, two of which were targeted exclusively in plastids, MORF2 and MORF9. MORF8 was a dual-localized protein which targeted to mitochondria and plastids. The remaining two MORFs, MORF1 and MORF3 were targeted to mitochondria. Because of the dual-localization of OsPGL1, we further investigated the physical interactions between OsPGL1 and these MORF proteins in pairs. Firstly, we performed yeast two-hybrid assay, data showed that OsPGL1 can solid interact with three MORFs, MORF2, MORF8 and MORF9 in yeast. However, no interaction was observed between OsPGL1 and other two MORFs as well as the negative control (Fig. 7A). Interestingly, the interaction between OsPGL1 and MORF9 was much more stronger than those of other MORFs. Moreover, the interactions between OsPGL1 and MORF2/8/9 were also observed when we switched the bait and prey in a yeast two-hybrid system (Fig. S6). Data displayed the stronger interaction between OsPGL1 and MORF9, compared with others. Next, we performed GST pull-down assays to validate the interactions *in vitro*, the three MORF proteins were fused with His tag and OsPGL1 was fused with GST tag for expression (Fig. S8A). Recombinant proteins were further verified by western blots (Fig. S8B) and subjected for pull-down assay in pairs. Trx-MORF2-His, Trx-MORF8-His and Trx-MORF9-His were all pulled down by GST-OsPGL1, respectively, which demonstrated that OsPGL1 can interact with these three MORF proteins directly (Fig. 7B).

**Figure 7.**
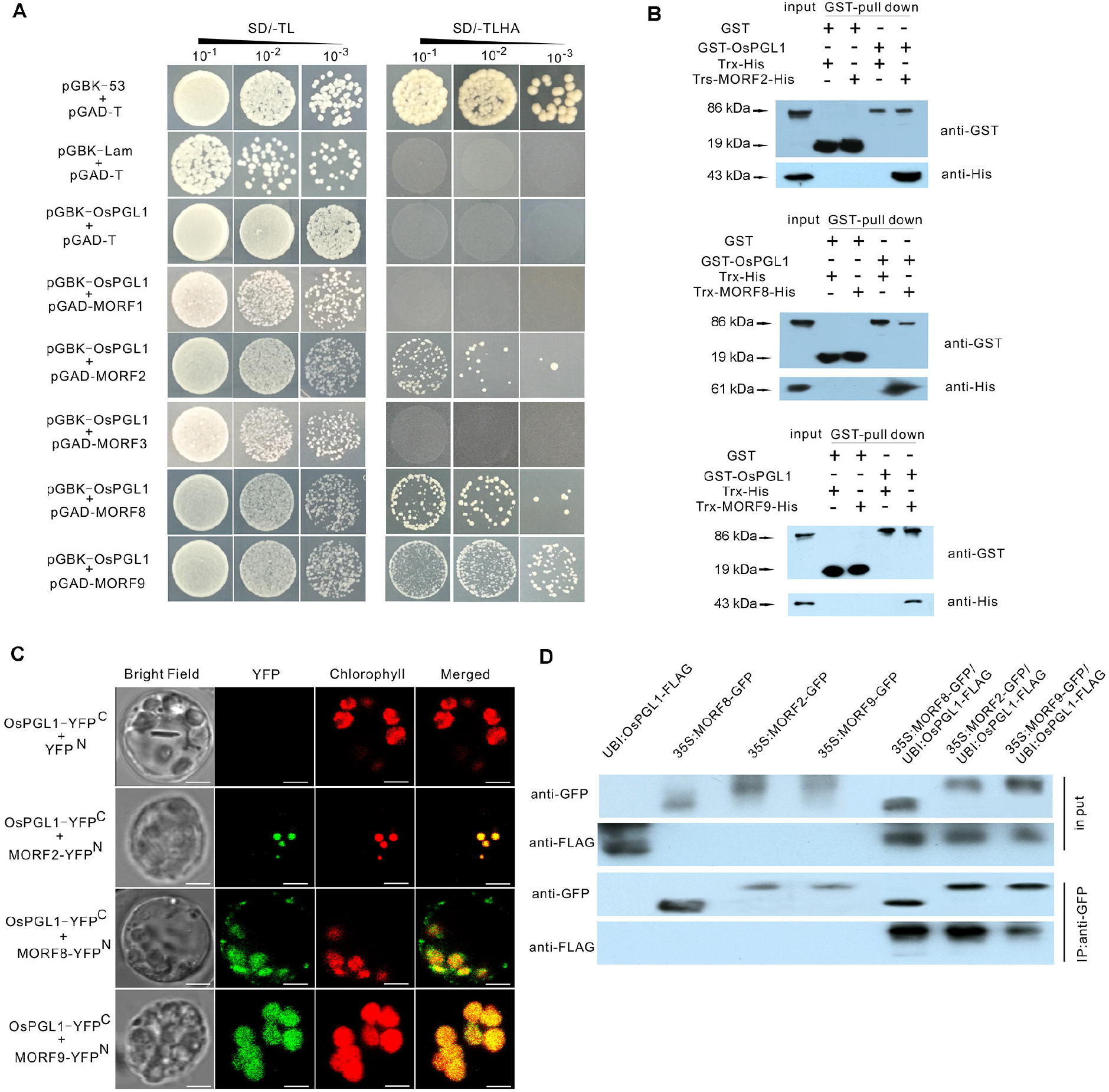
OsPGL1 directly interacts with MORF2/8/9 proteins. **(A)** Yeast-two hybrid assay, pGAD, GAL4 activation domain, used as prey vector, pGBK, GAL4 DNA binding domain, used as bait vector, SD/-TL and SD/-TLHA indicate SD/–Trp-Leu and SD/-Trp-Leu-His-Ade dropout plates, respectively. pGBK-53 and pGBK-lam was used as positive and negative control. Co-transcription of pGAD-T and pGBK-OsPGL1 used for self-activation detection. **(B)** GST Pull-down assay, the interaction between OsPGL1 and MORF2/8/9 proteins with detected by GST Pull-down assay, respectively. GST and Trx-His tag protein was used as control. The eluates were immunoblotted with anti-GST and anti-His antibodies, respectively. **(C)** Bimolecular fluorescence complementation assay showing that OsPGL1-YFP^C^ interacts with MORF2-YFP^N^, MORF8-YFP^N^ and MORF9-YFP^N^ to produce YFP fluorescence in the chloroplasts. Bar = 5μm. **(D)** Co-immunoprecipitation assay detection with anti-FLAG and anti-GFP antibodies, respectively.

To test the interactions *in vivo*, we performed bimolecular fluorescence complementation (BiFC) assays in rice protoplast. OsPGL1 and three MORFs were fused to the C-terminal and N-terminal of yellow fluorescent protein (YFP), respectively. Results showed that co-expression of OsPGL1-YFP^C^ and MORF2-YFP^N^/MORF8-YFP^N^/MORF9-YFP^N^ exhibited strong signals overlapped with chlorophyll, while the negative combination OsPGL1-YFP^C^ and YFP^N^ did not produce any detectable fluorescence signal (Fig. 7C). Interestingly, the signals from OsPGL1-YFP^C^ and MORF8-YFP^N^ were much more than chlorophyll, suggesting the interaction in mitochondria, which consistent with the dual-localization of OsPGL1 and MORF8.

Moreover, we generated the transgenic plants carrying UBI:OsPGL1-FLAG, 35S:MORF2-GFP, 35S:MORF8-GFP and 35S:MORF9-GFP, respectively. The protein crude extractions were incubated for co-immunoprecipitation assays with anti-GFP antibodies. Results confirmed the interactions between OsPGL1 and MORFs (Fig. 7D). Taken together, the results solidly demonstrated that OsPGL1 interacted with MORF2, MORF8 and MORF9 *in vitro* and *in vivo*.

### *Ospgl1* exhibited defective in photosynthetic complex

To make clear whether the photosynthetic and respiratory complex were impaired in *Ospgl1*, we detected the proteins involved in photosynthesis and electronic transfer chains pathway. Firstly, we examined NdhD in WT, *Ospgl1* mutants and the complementation line. Results showed greatly reduced accumulation of NdhD, suggesting the loss of editing at *ndhD-878* generated an unstable NdhD in chloroplast (Fig. 8A). Meanwhile, the accumulation of the photosystem I (PSI) subunits PsaA, PSII subunits PsbA, cytochrome b6f (PetA), chloroplast ATP synthase subunit AtpA, light harvesting complex of PSI (Lhca2) and light harvesting complex of PSII (Lhcb2) were also examined. All of these proteins were dramatically decreased in both two mutant lines (Fig. 8A). As expected, the levels of NdhD and other subunits in photosynthetic complex were recovered in the transgenic complementation line (Fig. 8A).

**Figure 8.**
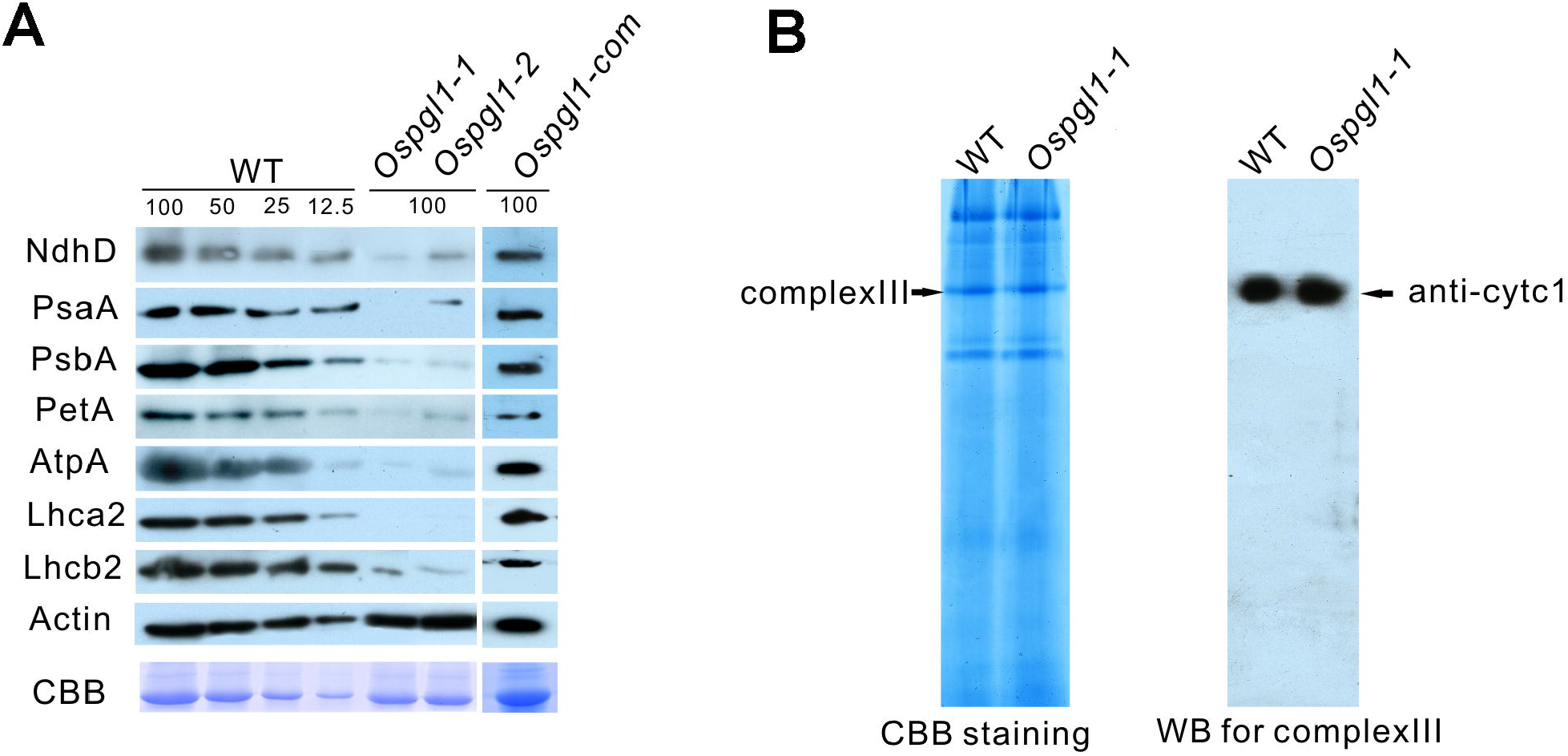
Immunoblotting analysis of the subunits of photosynthetic complexes and respiratory complex III. **(A)** Immunoblot analysis of the subunits of photosynthetic complex in WT, *Ospgl1* mutants, and complementation line at 4-leaf stage. Actin served as reference antibody, CBB staining indicated the loading control. Lanes were loaded with a series of dilutions as indicated. **(B)** The level of major protein of mitochondrial respiratory complex III analysis by BN-PAGE (left) and immunoblot (right). Cytc1 was used as a representative subunit of complex III.

We next examined the proteins involved in mitochondrial electron transport chain pathway. Mitochondrial complexes isolated from calli of *Ospgl1-1* and WT were separated on blue-native gel. No obvious changes were observed in all of the complexes via coomassie blue staining, implying *Ospgl1* did not compromise the function of mitochondria (Fig. 8B). Although the loss of RNA editing at *ccmFc-543* did not change the amino acid, we detected mitochondrial complex III as well, since CcmFc is a subunit of complex III. Protein immunoblotting with antibodies anti-CytC1 showed no difference at the amount of complex III in *Ospgl1-1* compared with that in WT (Fig. 8B), consistent with the undistinguishable morphological structure of mitochondria between *Ospgl1* and WT. Taken together, these results indicated photosynthetic complex was impaired while the respiratory complex was not affected in *Ospgl1* mutants.

### Altered expression of chloroplast development related genes in *Ospgl1*

The development of chloroplast in plant is related to the coordinated expression of both chloroplast and nuclear genes. Therefore, we first examined the transcript level of the chloroplast-encoded genes at 5-leaf stage of *Ospgl1* and WT plants, most of which are mediated by two types of RNA polymerase: plastid-encoded polymerase (PEP) and nuclear-encoded polymerase (NEP). Results showed that the expression levels of PEP-dependent genes (*rbcL*, *psaA*, *psbA*, and *petB*) were reduced in *Ospgl1*, which is consistent with results of the detection of protein level. Whereas, the NEP-dependent genes: RNA polymerase (*rpoA* and *rpoB*) were activated in *Ospgl1* (Fig. 9A). Results implied that some retrograde signals from chloroplast might activate NEP-dependent genes transcription for compensation in the development of chloroplast.

**Figure 9.**
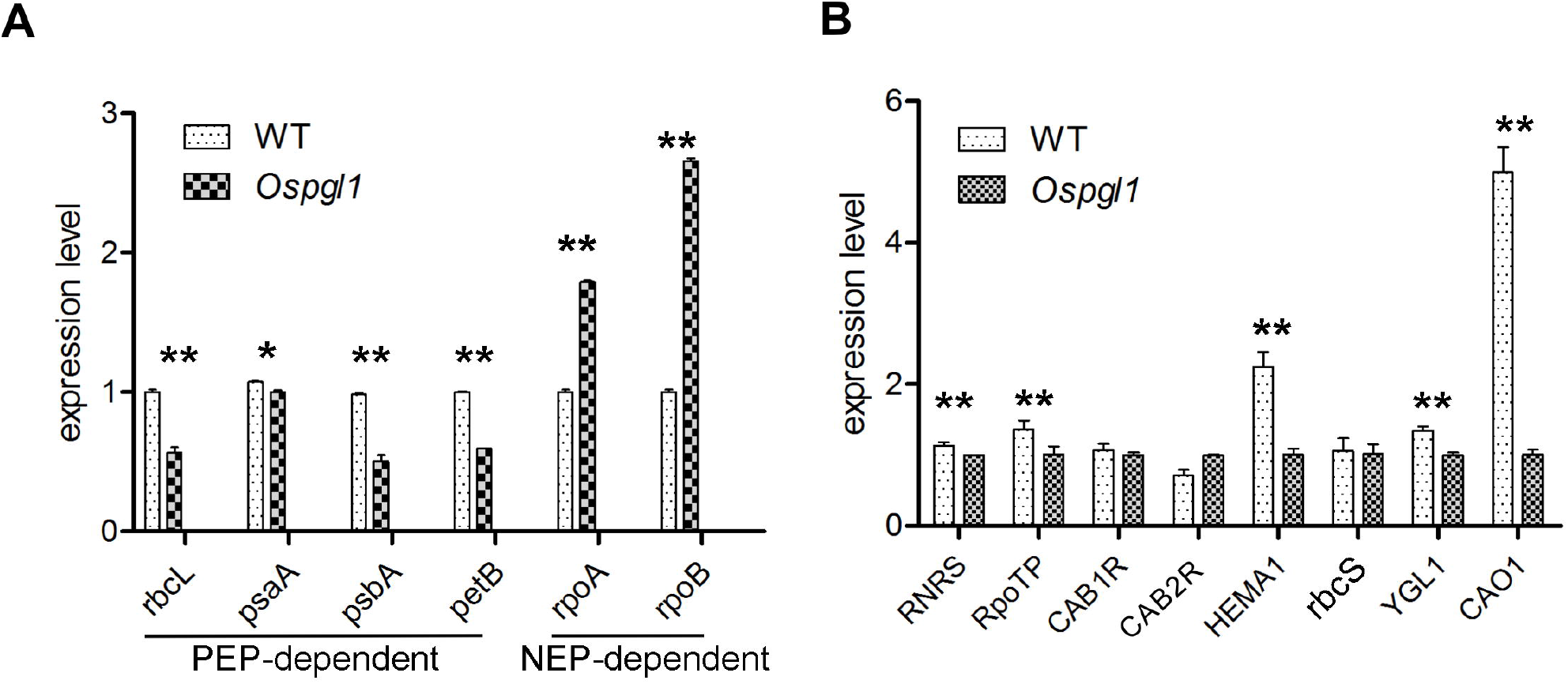
Expression Analysis of chloroplast development related genes. **(A)** Real time RT-PCR examination of PEP-dependent genes (rbcL, *psaA, psbA*, and *petB*) and NEP-dependent genes (*rpoA* and *rpoB*) in WT and *Ospgl1* at 5-leaf stage. Error bars represent the SD. (Student’s t-test: * P< 0.05, ** P< 0.01) **(B)** Real time RT-PCR examination of chloroplast development and photosynthesis related genes in WT and *Ospgl1* at 5-leaf stage. Error bars represent the SD. (Student’s t-test: ** P< 0.01)

Besides chloroplast-encoded genes, we also investigated the expression of some nuclear-encoded genes related to chloroplast development and photosynthesis in *Ospgl1* and WT plants, including *RNRS* (encoding the large subunit of RNR), *RpoTp* (encoding NEP core subunits), *CAB1R* and *CAB2R* (light-harvesting Chl a/b-binding protein of PSII), *HEMA1* (encoding a glutamyl-tRNA reductase), *rbcS* (encoding a Rubisco small subunit), *YGL1* (encoding a chlorophyll synthetase) and *CAO1* (encoding chlorophyll A oxygenase1). qRT–PCR analysis showed that the expression level of these genes was reduced in the *Ospgl1* mutant (Fig. 9B). Taken together, these data indicated that *OsPGL1* plays an important role in regulating the chloroplast development and photosynthesis.

### OsPGL1 recognizes the target RNA sequence

PPR protein binds target RNA via a modular recognition mechanism. To evaluate the conservation of the P, L, S motifs of OsPGL1, the sequence of each motif were analyzed. The alignments of these 9 motifs showed that residue Thr, Ala and Ser at position 6, Asn and Asp at position 1’ was conserved polar residues (Fig. 10A). More interestingly, residue Gly is also highly conserved at position 16, especially completely conserved at position 33 in P and L motifs (Fig. 10A). Data implied Gly might be essential for the function of PPR motif. To evaluate the match degree of OsPGL1 protein binding to its two target transcripts, we did a computational prediction. Alignments of the target sites of OsPGL1 showed comparatively high matches of PPR-RNA recognition basis with these two editing sites (Fig. 10B). The alignments also suggested the combinations between PPR motifs and its targeted nucleotides might be very flexible.

**Figure 10.**
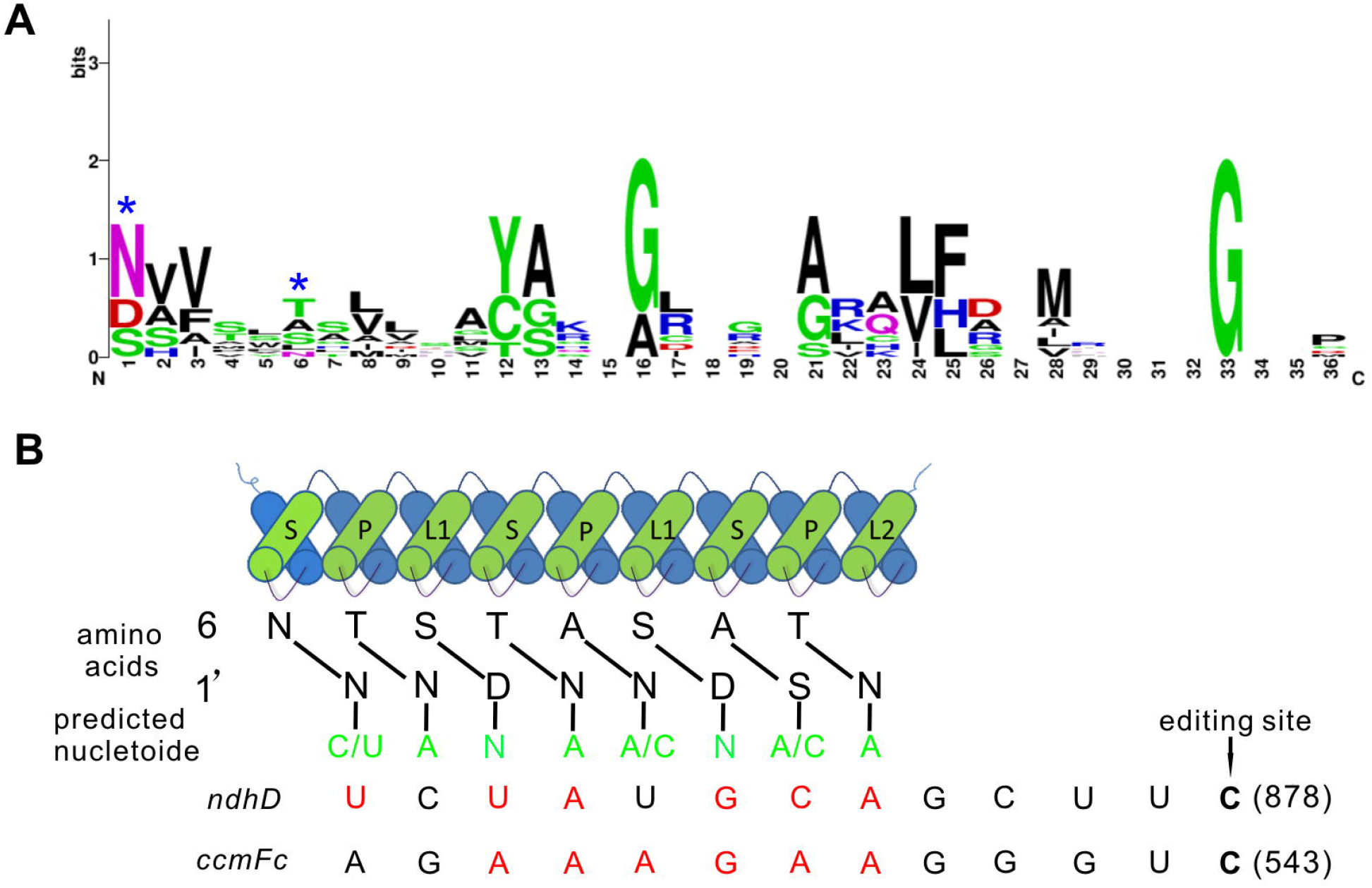
Alignment of PPR motifs of OsPGL1 with the cis-elements. **(A)** Sequence logo for PPR motifs in OsPGL1, the two positions that contribute to RNA binding specificity was indicated by stars. Sequence logos were constructed by Web-Logo. **(B)** Alignment of amino acid residues at position 6 and 1’ in each neighboring PPR motif of OsPGL1 with the putative cis-elements surrounding the editing sites. Nucleotides matching the PPR-RNA recognition combination were marked in red.

## Discussion

### OsPGL1 is a dual-localization PPR protein for RNA editing

PPR proteins were confirmed as a trans-acting factor for organelle RNA and acted as site specific RNA binding proteins (Okuda *et al.*, 2006). Most PPRs were involved in organelle RNA editing either mitochondrion or chloroplasts specifically, few of them are dual-localization. SOAR1 is a cytosol-nucleus dual-localized pentatricopeptide repeat (PPR) protein, which acting downstream of CHLH/ABAR and upstream of a nuclear ABA-responsive bZIP transcription factor ABI5 (Jiang *et al.*, 2015). PNM1 is dual localized to mitochondria and nuclei in Arabidopsis, which was only associated with polysomes and played a role in translation in mitochondria, and interacted with TCP8 in the nucleus (Hammani *et al.*, 2011). Both PPR2263 and MEF29 dually targeted to mitochondria and chloroplasts, and are required for RNA editing in maize and Arabidopsis, respectively (Sosso *et al.*, 2012). Here, we firstly reported a novel dual-localization PPR protein to mitochondria and chloroplasts in rice. Our data showed OsPGL1 is also required for RNA editing, and binds to target RNA directly. Both *ndhD-878* (chloroplast) and ccmFc-543 (mitochondrion) were completely edited in WT, even though the editing of ccmFc-543 (mitochondrion) is synonymous. Loss of function of OsPGL1 leads to the editing efficiency decrease to zero, and resulted in the pale green leaf phenotype. The dual-localization of OsPGL1 suggested the signal peptide sequence of OsPGL1 could be applied for trans-locating an artificial protein for mitochondria and chloroplasts synchronously in future.

### OsPGL1 recognizes target RNAs and functions with MORF proteins

The PPR motif consists of two anti-parallel α-helixes, the binding specificity depend on the first α-helix (Barkan *et al.*, 2012). Bioinformatic and structural analysis indicated that three positions of amino acids distributed in two adiacent PPR repeats were of great importance in recognizing its target RNA base (Takenaka *et al.*, 2013a). The alignment of OsPGL1 showed that high conservation of these positions in monocots (Fig. S5), implying the conserved function of those orthologs genes in RNA editing of *ndhD* and *ccmFc* in monocots. OsPGL1 was a DYW subgroup PPR protein with 3 P, 3 L and 3S motifs. More interesting finding is that the highly conservation of Gly and Ala at position 16, and completely conservation of Gly at position 33 in P and L motifs, which could be important for the function of PPR proteins. In this study, the RNA recognition also follows the rules that the editing site is located at downstream of binding sites, and the interval space is 4 nucleotides. In a word, such modular mode can help us to deeply understand the RNA recognition of PPR proteins and its application in future.

Non-PPR editing factors RIPs/MORFs, ORRM, OZ, and PPO1 have been identified as components of the plant RNA editosome, which was required for RNA editing (Sun *et al.*, 2016). In Arabidopsis plastids, both of the two plastid-localized members, MORF2 and MORF9 were required for RNA editing for mostly sites (Takenaka *et al.*, 2012). Recent study showed the RNA-binding activity of an artificial (PLS)_3_PPR could be sharply increased upon MORF9 binding, suggested the interaction between PPR and MORF9 could be more vital than that of others (Yan et al., 2017). In Arabidopsis, CLB19 interacted with MORF2, PDM2 interacted with MORF2/9, all of PDM1/SEL1, PPO1 and ORRM6 interacted with MORF2/8/9 (Du *et al.*, 2017; Hackett *et al.*, 2017; Ramos-Vega *et al.*, 2015; Zhang *et al.*, 2014; Zhang *et al.*, 2015). All these data indicated the editosome in plant is quite complicated. Nevertheless, MORF proteins were rarely reported in rice. Recently, WSP1, a sequence similarity with MORF protein was identified in rice, which was involved in RNA editing and splicing of several plastid genes (Zhang *et al.*, 2017). In our study, we confirmed that OsPGL1 can also interact with MORF2, MORF8 and MORF9 *in vitro* and *in vivo*. The interaction between OsPGL1 and MORF9 was significant stronger than that of others, implying the RNA binding activities of OsPGL1 could be enhanced by cooperation with MORF9. According to our data, we believe that OsPGL1 conducts the organelle RNA editing via an editosome coupled with MORF proteins. Other factors or subunits of editosome will be further explored based on our transgenic plants carrying UBI: OsPGL1-FLAG, 35S:MORF2-GFP, 35S:MORF8-GFP and 35S:MORF9-GFP.

### OsPGL1 plays an important role in rice chloroplast development

Leaf is an important tissue for plant. In this study, we constructed CRISPR/Cas9 knockout mutants of PPR gene *OsPGL1, Ospgl1-1* and *Ospgl1-2*, both of which exhibited pale green leaves during the whole vegetative stages. Further investigation revealed the chloroplast development and photosynthesis were defective on RNA and protein level in the mutant. NEP-dependent genes were activated in mutant implied some unknown retrograde signals from chloroplast were involved in compensation effects in the development of chloroplast. Complementation lines could rescue this aberrant phenotype, suggested that the base deletion mutation in *OsPGL1* gene was responsible for the PGL phenotype. OsPGL1 regulated chloroplast development by organelle RNA editing of *ndhD*. The highly conserved leucine at NdhD-293 is important for the structure or function of NdhD. We proposed the loss of RNA editing might lead to unstable NdhD in chloroplast, based on the results of protein immunoblotting, which showed the amount of photosynthetic complex subunits were dramatically decreased in mutants. Results suggested that PPR genes play a vital role in regulating chloroplast development via RNA editing in plant.

## Acknowledgements

We thank Yaoguang Liu laboratory for providing us with pYLCRISPR/Cas9 vectors.

No conflict of interest declared

## Author contributions

H.X., J.Hu. and Y.Z. designed the study. H.X. and Q.Z. contributed to constructed and *Ospgl1-1, Ospgl1-2* mutants and RNAi lines. H.X. and Y.X. carried out most experiments, H.X. and Q.Z. conducted SEM and TEM, F.Z. and C.N. performed expression and purification of recombinant proteins, J.Huang. contributed to field managements, H.X. and J.Hu. wrote the manuscript with feedback from all authors.

## Supplemental data

**Figure S1**. Phenotype of the *Ospgl1* and WT in a paddy field.

**Figure S2**. Phenotype of WT and RNAi line.

**Figure S3**. Schematic structural sequence of OsPGL1.

**Figure S4**. Sequence alignment of *OsPGL1* with its orthologs in various plants.

**Figure S5**. Conservation of the amino acid at 6 and 1’ position of each PPR motif.

**Figure S6**. Interaction of OsPGL1 with MORF2/MORF8/MORF9 by yeast two-hybrid assay.

**Figure S7**. Expression and purification of GST-OsPGL1^46-604^.

**Figure S8**. Expression and purification of Trx-MORF2-His, Trx-MORF8-His and Trx-MORF9-His.

**Table S1**. Design of target adaptor for CRISPR/Cas9 system.

**Table S2**. Primers used for qRT-PCR, vector construction and RNA editing.

## References

Barkan A, Rojas M, Fujii S, Yap A, Chong YS, Bond CS, Small I. 2012. A combinatorial amino acid code for RNA recognition by pentatricopeptide repeat proteins. PLoS Genet 8, e1002910.

Bentolila S, Babina AM, Germain A, Hanson MR. 2014. Quantitative trait locus mapping identifies REME2, a PPR-DYW protein required for editing of specific C targets in Arabidopsis mitochondria. RNA Biol 10, 1520–1525.

Bentolila S, Heller WP, Sun T, Babina AM, Friso G, van Wijk KJ, Hanson MR. 2012. RIP1, a member of an Arabidopsis protein family, interacts with the protein RARE1 and broadly affects RNA editing. Proc Natl Acad Sci US A 109, E1453–1461.

Chateigner-Boutin AL, Small I. 2010. Plant RNA editing. RNA Biol 7, 213–219.

Cheng S, Gutmann B, Zhong X, Ye Y, Fisher MF, Bai F, Castleden I, Song Y, Song B, Huang J, Liu X, Xu X, Lim BL, Bond CS, Yiu SM, Small I. 2016. Redefining the structural motifs that determine RNA binding and RNA editing by pentatricopeptide repeat proteins in land plants. Plant J 85, 532–547.

Covello PS, Gray MW. 1989. RNA editing in plant mitochondria. Nature 341, 662–666.

Du L, Zhang J, Qu S, Zhao Y, Su B, Lv X, Li R, Wan Y, Xiao J. 2017. The Pentratricopeptide Repeat Protein Pigment-Defective Mutant2 is Involved in the Regulation of Chloroplast Development and Chloroplast Gene Expression in Arabidopsis. Plant Cell Physiol 58, 747–759.

Hackett JB, Shi X, Kobylarz AT, Lucas MK, Wessendorf RL, Hines KM, Bentolila S, Hanson MR, Lu Y. 2017. An Organelle RNA Recognition Motif Protein Is Required for Photosystem II Subunit psbF Transcript Editing. Plant Physiol 173, 2278–2293.

Hammani K, Gobert A, Hleibieh K, Choulier L, Small I, Giege P. 2011. An Arabidopsis dual-localized pentatricopeptide repeat protein interacts with nuclear proteins involved in gene expression regulation. Plant Cell 23, 730–740.

Hess GT, Fresard L, Han K, Lee CH, Li A, Cimprich KA, Montgomery SB, Bassik MC. 2016. Directed evolution using dCas9-targeted somatic hypermutation in mammalian cells. Nat Methods 13, 1036–1042.

Hu J, Wang K, Huang W, Liu G, Gao Y, Wang J, Huang Q, Ji Y, Qin X, Wan L, Zhu R, Li S, Yang D, Zhu Y. 2012. The rice pentatricopeptide repeat protein RF5 restores fertility in Hong-Lian cytoplasmic male-sterile lines via a complex with the glycine-rich protein GRP162. Plant Cell 24, 109–122.

Jiang SC, Mei C, Liang S, Yu YT, Lu K, Wu Z, Wang XF, Zhang DP. 2015. Crucial roles of the pentatricopeptide repeat protein SOAR1 in Arabidopsis response to drought, salt and cold stresses. Plant Mol Biol 88, 369–385.

Kim D, Lim K, Kim ST, Yoon SH, Kim K, Ryu SM, Kim JS. 2017. Genome-wide target specificities of CRISPR RNA-guided programmable deaminases. Nat Biotechnol 35, 475–480.

Kim U, Wang Y, Sanford T, Zeng Y, Nishikura K. 1994. Molecular cloning of cDNA for double-stranded RNA adenosine deaminase, a candidate enzyme for nuclear RNA editing. Proc Natl Acad Sci U S A 91, 11457–11461.

Liu G, Tian H, Huang YQ, Hu J, Ji YX, Li SQ, Feng YQ, Guo L, Zhu YG. 2012. Alterations of mitochondrial protein assembly and jasmonic acid biosynthesis pathway in Honglian (HL)-type cytoplasmic male sterility rice. J Biol Chem 287, 40051–40060.

Lu Y, Zhu JK. 2017. Precise Editing of a Target Base in the Rice Genome Using a Modified CRISPR/Cas9 System. Mol Plant 10, 523–525.

Lurin C, Andres C, Aubourg S, Bellaoui M, Bitton F, Bruyere C, Caboche M, Debast C, Gualberto J, Hoffmann B, Lecharny A, Le Ret M, Martin-Magniette ML, Mireau H, Peeters N, Renou JP, Szurek B, Taconnat L, Small I. 2004. Genome-wide analysis of Arabidopsis pentatricopeptide repeat proteins reveals their essential role in organelle biogenesis. Plant Cell 16, 2089–2103.

Ma Y, Zhang J, Yin W, Zhang Z, Song Y, Chang X. 2016. Targeted AID-mediated mutagenesis (TAM) enables efficient genomic diversification in mammalian cells. Nat Methods 13, 1029–1035.

McDougall WM, Okany C, Smith HC. 2011. Deaminase activity on single-stranded DNA (ssDNA) occurs in vitro when APOBEC3G cytidine deaminase forms homotetramers and higher-order complexes. J Biol Chem 286, 30655–30661.

Mehta A, Driscoll DM. 2002. Identification of domains in apobec-1 complementation factor required for RNA binding and apolipoprotein-B mRNA editing. RNA 8, 69–82.

Melcher T, Maas S, Herb A, Sprengel R, Seeburg PH, Higuchi M. 1996. A mammalian RNA editing enzyme. Nature 379, 460–464.

Okuda K, Nakamura T, Sugita M, Shimizu T, Shikanai T. 2006. A pentatricopeptide repeat protein is a site recognition factor in chloroplast RNA editing. J Biol Chem 281, 37661–37667.

Ramos-Vega M, Guevara-Garcia A, Llamas E, Sanchez-Leon N, Olmedo-Monfil V, Vielle-Calzada JP, Leon P. 2015. Functional analysis of the Arabidopsis thaliana CHLOROPLAST BIOGENESIS 19 pentatricopeptide repeat editing protein. New Phytol 208, 430–441.

Salone V, Rudinger M, Polsakiewicz M, Hoffmann B, Groth-Malonek M, Szurek B, Small I, Knoop V, Lurin C. 2007. A hypothesis on the identification of the editing enzyme in plant organelles. FEBS Lett 581, 4132–4138.

Schmitz-Linneweber C, Small I. 2008. Pentatricopeptide repeat proteins: a socket set for organelle gene expression. Trends Plant Sci 13, 663–670.

Small ID, Peeters N. 2000. The PPR motif - a TPR-related motif prevalent in plant organellar proteins. Trends Biochem Sci 25, 46–47.

Sosso D, Mbelo S, Vernoud V, Gendrot G, Dedieu A, Chambrier P, Dauzat M, Heurtevin L, Guyon V, Takenaka M, Rogowsky PM. 2012. PPR2263, a DYW-Subgroup Pentatricopeptide repeat protein, is required for mitochondrial nad5 and cob transcript editing, mitochondrion biogenesis, and maize growth. Plant Cell 24, 676–691.

Sun T, Bentolila S, Hanson MR. 2016. The Unexpected Diversity of Plant Organelle RNA Editosomes. Trends Plant Sci 21, 962–973.

Takenaka M, Zehrmann A, Brennicke A, Graichen K. 2013a. Improved computational target site prediction for pentatricopeptide repeat RNA editing factors. PLoS One 8, e65343.

Takenaka M, Zehrmann A, Verbitskiy D, Hartel B, Brennicke A. 2013b. RNA editing in plants and its evolution. Annu Rev Genet 47, 335–352.

Takenaka M, Zehrmann A, Verbitskiy D, Kugelmann M, Hartel B, Brennicke A. 2012. Multiple organellar RNA editing factor (MORF) family proteins are required for RNA editing in mitochondria and plastids of plants. Proc Natl Acad Sci US A 109, 5104–5109.

Teng B, Burant CF, Davidson NO. 1993. Molecular cloning of an apolipoprotein B messenger RNA editing protein. Science 260, 1816–1819.

Yin P, Li Q, Yan C, Liu Y, Liu J, Yu F, Wang Z, Long J, He J, Wang HW, Wang J, Zhu JK, Shi Y, Yan N. 2013. Structural basis for the modular recognition of single-stranded RNA by PPR proteins. Nature 504, 168–171.

Yu C, Wang L, Chen C, He C, Hu J, Zhu Y, Huang W. 2014. Protoplast: a more efficient system to study nucleo-cytoplasmic interactions. Biochem Biophys Res Commun 450, 1575–1580.

Zhang F, Tang W, Hedtke B, Zhong L, Liu L, Peng L, Lu C, Grimm B, Lin R. 2014. Tetrapyrrole biosynthetic enzyme protoporphyrinogen IX oxidase 1 is required for plastid RNA editing. Proc Natl Acad Sci US A 111, 2023–2028.

Zhang HD, Cui YL, Huang C, Yin QQ, Qin XM, Xu T, He XF, Zhang Y, Li ZR, Yang ZN. 2015. PPR protein PDM1/SEL1 is involved in RNA editing and splicing of plastid genes in Arabidopsis thaliana. Photosynth Res 126, 311–321.

Zhang Z, Cui X, Wang Y, Wu J, Gu X, Lu T. 2017. The RNA Editing Factor WSP1 Is Essential for Chloroplast Development in Rice. Mol Plant 10, 86–98.

Zhou S, Wang Y, Li W, Zhao Z, Ren Y, Wang Y, Gu S, Lin Q, Wang D, Jiang L, Su N, Zhang X, Liu L, Cheng Z, Lei C, Wang J, Guo X, Wu F, Ikehashi H, Wang H, Wan J. 2011. Pollen semi-sterility1 encodes a kinesin-1-like protein important for male meiosis, anther dehiscence, and fertility in rice. Plant Cell 23, 111–129.

Zong Y, Wang Y, Li C, Zhang R, Chen K, Ran Y, Qiu JL, Wang D, Gao C. 2017. Precise base editing in rice, wheat and maize with a Cas9-cytidine deaminase fusion. Nat Biotechnol 35, 438–440.

